# Galvanic vestibular stimulation alters the sense of upright

**DOI:** 10.1101/2025.07.29.667402

**Authors:** Sofia Müller-Wöhrstein, Hans-Otto Karnath

## Abstract

Given the vestibular system’s important role in the perception of upright, we investigated the possible effects of galvanic vestibular stimulation (GVS) on the perception of one’s own upright body orientation in relation to gravity, the so-called ‘Subjective Postural Vertical (SPV)’. Two groups of healthy participants with an average age of 25.4 years and 64.5 years respectively, each consisting of 28 healthy participants, sat (blindfolded) on a tilting chair. The subjects’ feeling of being upright was tested under three different conditions of GVS: right-sided anodal stimulation, left-sided anodal stimulation, and sham stimulation. Our findings revealed that right-sided anodal GVS significantly altered the SPV in both age groups, whereas left-sided anodal GVS did not. The observed effect of GVS on perceived upright body posture was numerically small (up to 0.87° on average) and not due to a loss of sensitivity to the perception of body verticality. The unexpected asymmetry of the behavioral effects of GVS could be related to the known right hemispheric asymmetry of cortical activation in vestibular projection areas, which would need to be further clarified in future studies.

## Introduction

Neurological patients with pusher syndrome are characterized by a profound disorder of body orientation in space (Davies, 1985; Karnath, 2007). They perceive themselves as upright, although in reality they are tilted on average 18° to the side of their stroke (Bergmann et al., 2016; Karnath et al., 2000). In terms of compensation, it would therefore be valuable to use a method that could influence pusher patients’ erroneous perception of upright body orientation in relation to gravity. One modality for such manipulation could be the vestibular system. It plays a central role in the perception of upright body orientation, particularly in relation to gravity (Cohen et al., 2012b; Day & Fitzpatrick, 2005). Galvanic vestibular stimulation (GVS) is an non-invasive, electrical stimulation method that targets the vestibular system. Its detailed neurophysiological mechanisms are still a subject of debate. Some researchers argue that GVS primarily stimulates the otolith organs (Cohen et al., 2012a; Cohen et al., 2012b), while others debate that it primarily affects the semicircular canals (Curthoys & MacDougall, 2012). Also at the behavioral level, some questions are still being discussed. It is known that the main behavioral effect of GVS is directed to the side of the anode (Balter et al., 2004; Fitzpatrick & Day, 2004), eliciting, among others, otolith-related behavioral responses, such as a sense of roll and postural responses (Cohen et al., 2012b), especially in the absence of visual input (Curthoys & MacDougall, 2012). Also, it is known that GVS provokes a tilt of the so-called *Subjective Visual Vertical* (*SVV*; Mars et al., 2005; Tardy-Gervet & Séverac-Cauquil, 1998; Zink et al., 1998), that is a subject’s perception of the visual environment with respect to one’s own body orientation. But does GVS also alter the perception of one’s own upright body orientation in relation to gravity, that is the so-called *Subjective Postural Vertical* (*SPV*)? This latter perception in particular would be the one to influence in stroke patients with pusher syndrome.

The SPV is measured by having participants (blindfolded) sit or stand in tilting devices and align their body according to their subjective perception of the earth’s verticality (Dakin & Rosenberg, 2018). Additionally, the precision of one’s own upright body orientation in relation to gravity can be measured by using the values at entry and exit from perceived verticality, the so-called ‘sector width’ (Bisdorff et al., 1996). Apart from the fact that the application of GVS induces a postural sway when standing (Latt et al., 2003), the question of whether or not GVS alters the sense of upright in healthy subjects has not yielded clear results so far. Bisdorff et al. (1996) applied GVS during the measurement of the SPV in young adults. They found a larger sector width during GVS, that is they found a lower precision with which a participant indicated verticality, but no effect of GVS on the SPV itself. Volkening et al. (2014) tested the influence of GVS on various measures of verticality perception, including the SPV. As Bisdorff et al. (1996), the authors found no significant effects of GVS on the SPV. Yet, other studies mention an effect of GVS on body verticality, but without giving exact values (Mars et al., 2005) or describing the sensation of a change in gravity only qualitatively (Nguyen et al., 2022). It is essential to note that almost all of these earlier studies had a very small sample with a maximum of ten participants per study, especially the two studies with explicit measurements of the SPV. This may be too small to detect or to disprove possible effects of GVS on body orientation.

Another unresolved aspect is the possible influence of age on the perception of one’s own upright body orientation in relation to gravity under GVS. We know that the vestibular system of older people is less sensitive, and its input is less weighted (Alberts et al., 2019; Faraldo-García et al., 2012; Nestmann et al., 2020). This could lead to greater uncertainty and lower accuracy in the measurement of the SPV, but no studies have yet been conducted to measure the SPV during GVS in a larger sample of healthy older adults. Considering that stroke patients who could potentially benefit from these results in the future are on average over 65 years old (Dai et al., 2022), a potential age effect is of particular importance. The present study thus compared the behavioral effects of GVS between younger and older subgroups in a large sample of 56 healthy participants to answer the question whether or not the perception of upright body posture in relation to gravity can be influenced by GVS.

## Methods and Materials

### Participants

Participants were recruited through in-house mailing lists for research participation. For the younger group, a total of 29 neurologically healthy individuals (18 females) between the age of 20 and 31 (*M* = 25.2; *SD* = 3.4) participated in the study. The older group consisted of also 29 individuals (17 females), aged between 56 and 81 (*M* = 65.0; *SD* = 6.4). Upon arrival at the lab, all participants were allocated a subject code to guarantee the pseudonymity of their data. All participants gave their written informed consent in accordance with the revised version of the Declaration of Helsinki. The study was also approved by the Ethics Committee of the Medical Faculty of the University of Tübingen, Germany (814/2021BO2). After completion of testing participants were compensated monetarily for their participation.

### Galvanic Vestibular Stimulation (GVS)

Bilateral bipolar GVS was applied by using a CE-certified constant direct current stimulator (NeuroConn DC-Stimulator; neuroConn, Ilmenau, Germany). Before applying the electrodes on the skin binaurally over left and right mastoids, we cleared the skin using skin preparation gel (Nuprep skin prep gel; Weaver and Company, Aurora, Colorado, USA). The electrodes were then placed with conductive adhesive paste to minimize skin impedance (Ten20 Conductive Neurodiagnostic Electrode Paste; Weaver and Company, Aurora, Colorado, USA). Experimental stimulation was applied with a single mode current of 1mA for 15 seconds, with a fade-in and fade-out of 2 seconds each. For the sham stimulation condition (see below) only fade-in and fade-out was carried out, but without the 15 seconds of actual stimulation, so that the participants had the same tingling or warm sensation as in the experimental stimulation conditions (see below).

### Experimental Procedure

After attaching the electrodes, participants were seated on an in-house built tilting chair (Fig. 1), which can be tilted to the left and right by the experimenter by manually turning a wheel. The degree of tilt is displayed to the experimenter on a scale attached to the back of the tilting chair. Participants’ eyes were occluded using an eye mask. Additionally, the participants were well padded in their seat so that they would get as minimal somatosensory feedback as possible from the tilts and potential pressure on arms, legs and/or the buttocks.

**Fig. 1.**
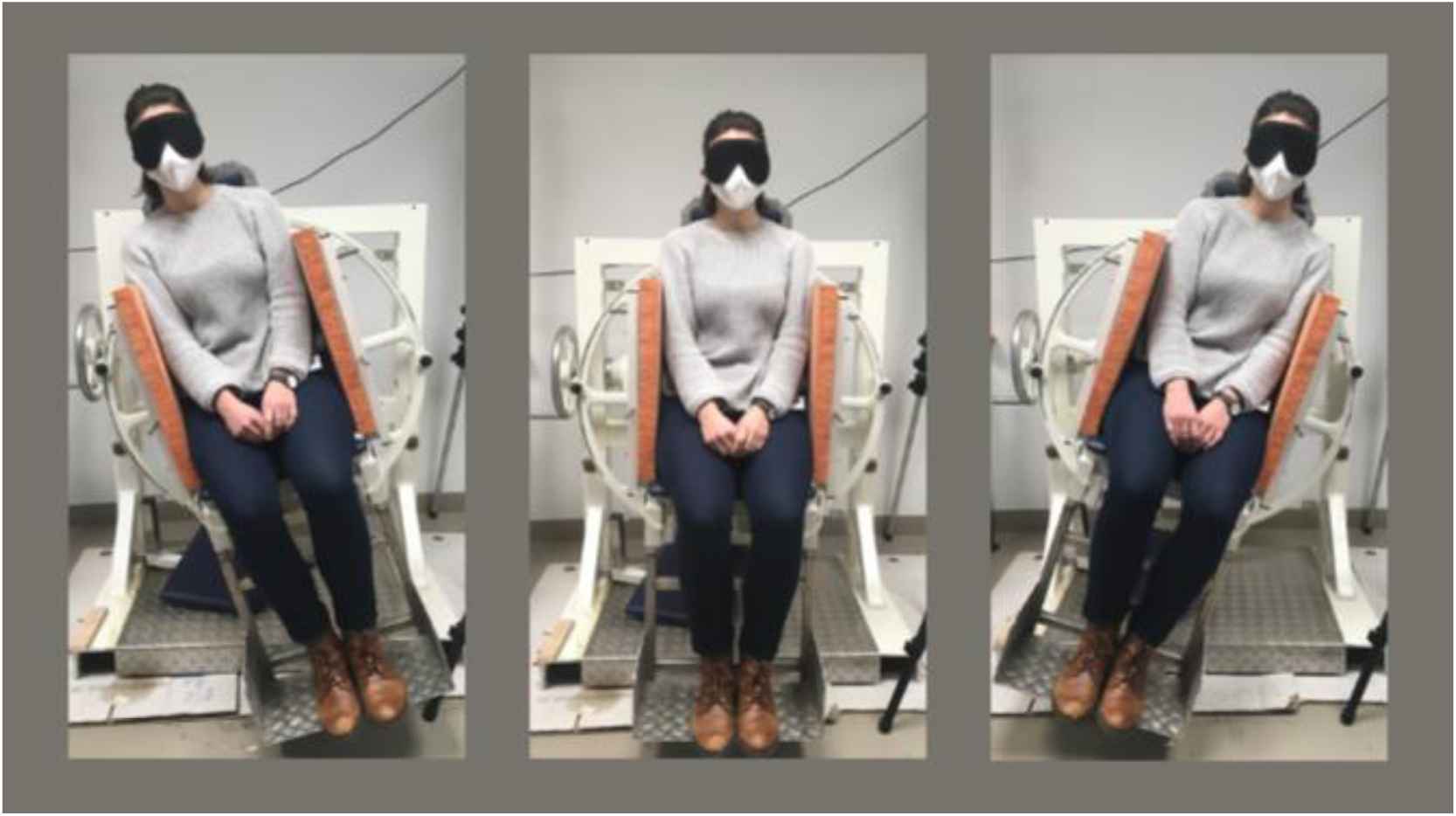
A photograph of one of the authors seated in the tilting chair. The tilting chair can be tilted to the left and right. During the experiment, participants wore an eye mask to exclude visual input and were well padded into the chair to minimize somatosensory feedback during tilts.

Tilts were performed in three experimental conditions: during anodal stimulation of the right mastoid (*‘anode right’*), during anodal stimulation of the left mastoid (*‘anode left’*) and during a sham stimulation (*‘sham’*). The participants were blinded to the three stimulation conditions. The starting condition as well as the order of conditions was randomized over all participants to prevent habituation effects to the vestibular stimulation.

For each condition eight tilts were performed (four starting from the right, four starting from the left). A total of 24 tilts were therefore carried out. A trial started with the tilting chair being driven to the starting tilt angle that was randomized for all eight tilts per condition but was the same for every participant (between 15° and 20° towards the left or right). Starting from this tilt position on the left or right, the tilting chair was turned slowly and regularly by the experimenter to the opposite side. The participants received the following instruction (before turning the tilting chair for the first time from its tarting tilt angle): “As soon as you have the feeling of being upright, please say *‘start’* and then say *‘stop’* as soon as you have the feeling that you are no longer upright.” The angle when *‘start’* was said was taken as the entry value into the subjective postural vertical, while the angle at *‘stop’* was considered the exit value from the SPV.

### Statistical Analyses

To calculate the mean SPV of a participant, the mean of the entry and exit values was taken and averaged over all trials per condition. In addition to the estimation of body verticality, we also determined the precision with which a participant indicates verticality. For the latter, we calculated the mean sector width per subject, that is the absolute difference between the entry and exit values and likewise averaged over all trials per condition. In this way, we obtained an averaged SPV value and an averaged sector width value for each participant in each condition. For all subsequent analyses, we excluded outliers, that is participants who showed a deviation of more than two SD from the group mean of the SPV in the sham condition (separately for either the young or the older group).

For the frequentist statistical analysis, we first ran two mixed ANOVA models with the three experimental conditions as the within-subject factor and age group as the between-subject factor. For the dependent variables, we used mean SPV in one model and mean sector width in the other. The analysis was performed using the *anova_test()* function from the *rstatix* package in *R* (Kassambara, 2023). If the main effects of the mixed ANOVA model were statistically significant, Bonferroni corrected post-hoc tests were performed to examine the effectsin detail. In addition, we also conducted the same analysis using a Bayesian approach to also obtain evidence for or against the null hypothesis (no difference between the three experimental conditions or age groups). Using the *anovaBF()* function from the *BayesFactor* package (Rouder et al., 2012), we built our mixed ANOVA models as previously described, including participant identifiers as random effects. In addition, instead of the default Monte Carlo integration for Bayes Factor estimation, we increased the number of iterations to 500000 to achieve a proportional error of <1%. All analyses were performed with *R Studio* (R version 4.4.0; Posit Software, 2024).

## Results

Experimental stimulation did not have to be terminated at the request of a participant in either the young or the older group, as no undesirable side effects occurred . In each of the two age groups, one participant had to be excluded as being an outlier, that is their SPV in the sham condition deviated more than 2 SD from the group mean. This left us with 28 young participants (17 females) between the age of 20 and 31 (*M* = 25.4; *SD* = 3.4) and 28 older participants (16 females) between the age of 56 and 78 (*M* = 64.5; *SD* = 5.7) with whom we performed the statistical analyses.

### Subjective Postural Vertical (SPV)

In the frequentist statistical model with SPV as the dependent variable, there was no significant interaction effect between age group and experimental condition (*F*(2,86) = 0.94, *p* = .376, *η_p_^2^* = .017), but a significant main effect for both the age group (*F*(1, 54) = 5.93, *p* = .018, *η_p_^2^* = .099) as well as the experimental condition (*F*(2, 86) = 6.47, *p* = .005, *η_p_^2^* = .107; see Fig. 2). The deviation from objective 0° earth-vertical body orientation was significantly higher in the older group (*M* = 0.54°, *SD* = 1.05°) than in the younger group (*M* = 0.07°, *SD* = 0.93°). Moreover, the deviation was greater with right anodal stimulation (*M* = 0.62°, *SD* = 1.18°) than with left anodal stimulation (*M* = 0.17°, *SD* = 0.86°) and sham stimulation (*M* = 0.12°, *SD* = 0.94°). Post-hoc tests with Bonferroni correction revealed that the difference between left and right anodal stimulation was statistically significant (*p* = 0.012) as well as the difference between right anodal stimulation and sham stimulation (*p* = 0.009). In contrast, the difference between left anodal stimulation and sham stimulation was not statistically significant (*p* = .770).

**Fig 2.**
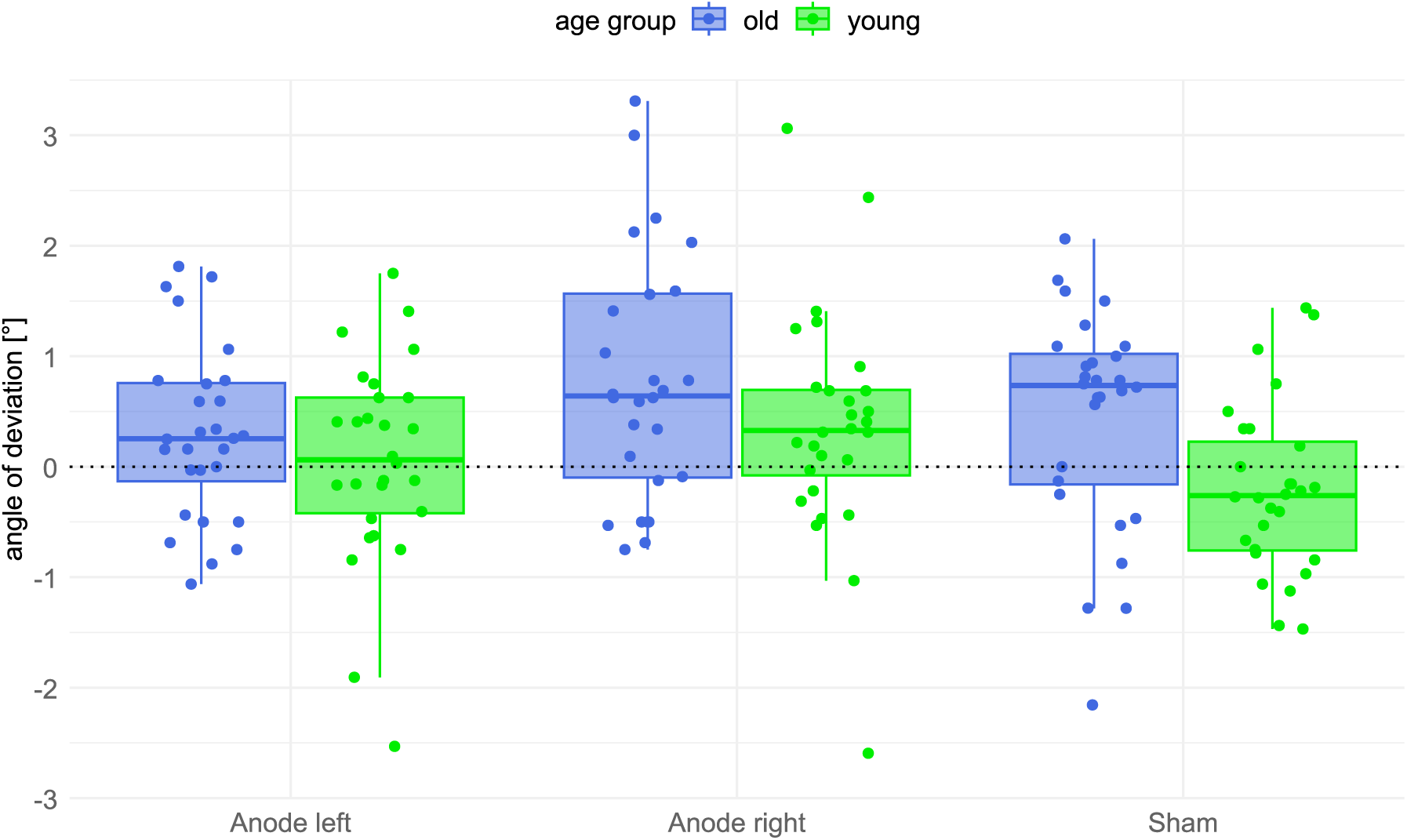
Mean subjective postural vertical (SPV) for each of the three conditions and both age groups. The boxplots show the deviation from objective 0° earth-vertical body orientation including the respective median and interquartile ranges. Dots show individual data points for each participant.

The Bayesian analysis showed a slightly different pattern than the frequentist approach. The ANOVA model including only the participant group was negligible (BF_10_ = 2.69). In contrast, the ANOVA model that only includes the experimental condition led to substantial evidence against the null hypothesis (BF_10_ = 14.13). For the ANOVA model that includes both factors, age group and experimental condition, there was strong evidence (BF_10_ = 39.13). Also, there was substantial evidence for the ANOVA model that also includes the interaction between the two effects (BF_10_ = 8.71), although considerably less than for the model without the interaction effect.

### Sector Width

In the frequentist statistical model with sector width as the dependent variable, there was no significant main effect for both the age group (*F*(1, 54) = 0.83, *p* = .368, *η_p_^2^* = .015) as well as the experimental condition (*F*(2, 90) = 0.21, *p* = .768, *η_p_^2^* = .004; see Fig. 3) There was also no significant interaction effect between age group and experimental condition (*F* (2,90) = 0.67, *p* = .488, *η_p_^2^* = .012). The Bayesian analysis corroborated these results. None of the four ANOVA models showed evidence against the null hypothesis (ANOVA model that included only the participant group [BF_10_ = 0.62], ANOVA model that included only the experimental condition [BF_10_ = 0.07], ANOVA model that included both factors, age group and experimental condition [BF_10_ = 0.05], ANOVA model that additionally included the interaction between the two effects [BF_10_ = 0.01]).

**Fig 3.**
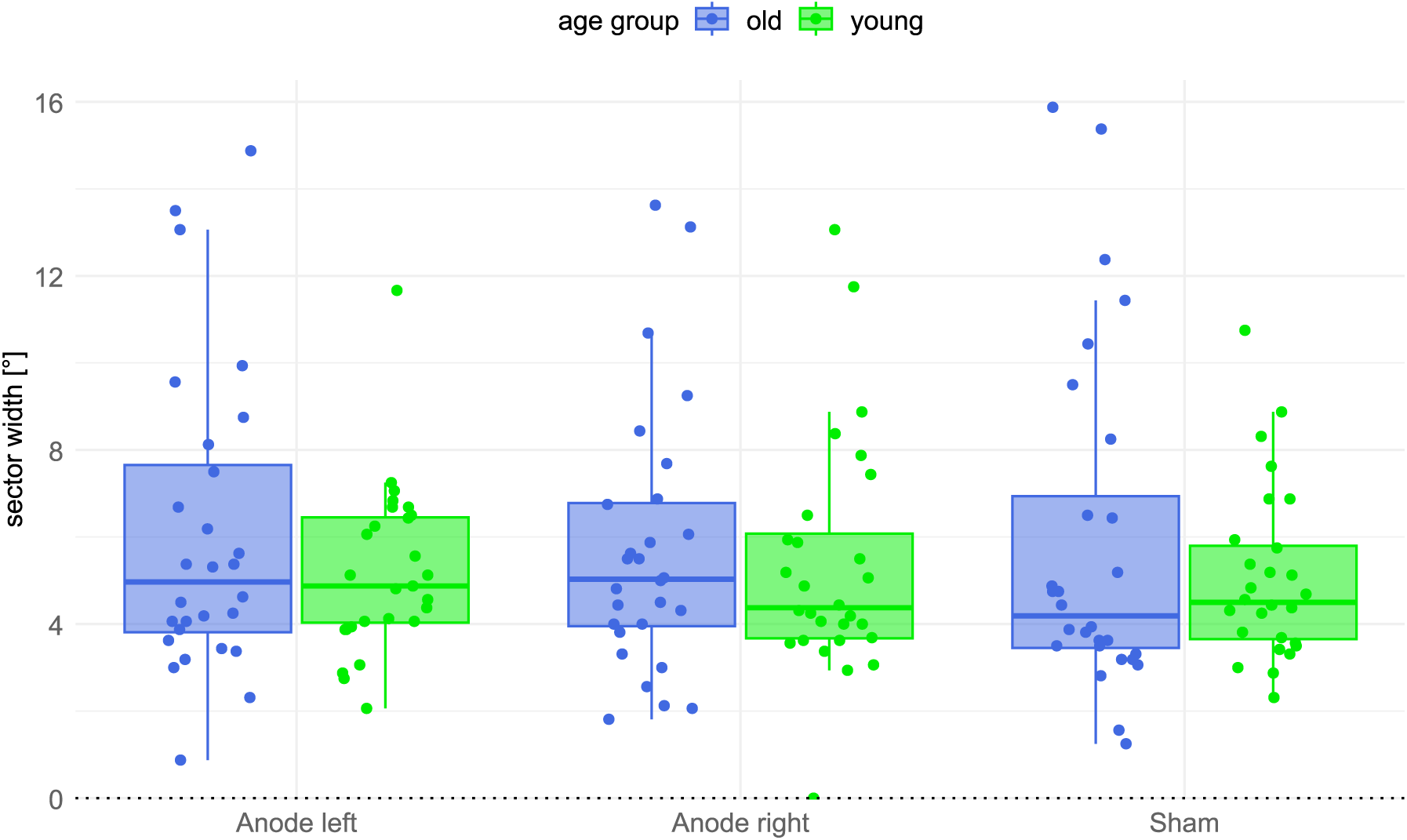
Sector width for each of the three conditions and both age groups. The boxplots show the absolute range between the entry and exit values into verticality including the respective median and interquartile ranges. Dots show individual data points for each participant.

### Correlation between Subjective Postural Vertical (SPV) and Sector Width

To test whether the mean SPV and the mean sector width are correlated, we calculated a Pearsońs correlation between the SPV value and the sector width for each of the three conditions, separately for the young and the old group. In the sham stimulation condition, the SPV and sector width were not significantly correlated, neither in the young group (r = -0.08, *p* = .691) nor in the older group (r = -0.22, *p* = .251). Similarly, the SPV and sector width were not significantly correlated in the right anodal stimulation condition, neither in the young group (r = 0.18, *p* = .355) nor in the older group (r = -0.17, *p* = .376), nor in the left anodal stimulation condition, again neither in the young group (r = -0.07, *p* = .725) nor in the older group (r = - 0.14, *p* = .483).

## Discussion

In a large sample of 56 healthy participants, the present study observed that GVS indeed had an effect on the SPV when anodal stimulation was performed on the right mastoid. This effect was supported by both frequentist as well as Bayesian statistics. Conversely, we found no such effect when anodal stimulation was performed on the left mastoid. Beyond, participants’ age appears to have an effect on the SPV; in all stimulation conditions we observed that older participants showed larger SPV deviations than younger participants. However, the influence of age seems to be small, as the Bayesian evidence was rather low, even though the frequentist approach yielded a significant result. In contrast to the SPV, the participants’ sensitivity for indicating upright body posture in relation to gravity, measured by the sector width, was neither influenced by GVS nor by age. In keeping with this, we found the SPV and the sector width as independent measures; the two variables did not show significant correlations. The latter results allow to conclude that the significant effect of GVS on perceived upright body posture is not due to a loss of sensitivity to the perception of body verticality.

The results obtained in our study differ in part from some of the earlier studies conducted on this topic (see introduction section). If one compares the ‘sector width’ values measured by Bisdorff et al. (1996) during right anodal stimulation, there is hardly any numerical difference to the present study (Bisdorff et al.: right anodal GVS 5.03°; present study: right anodal GVS 5.52°). Rather, a difference is found with regard to the control condition with which this value was compared to in the two studies. While Bisdorff et al. (1996) compared the right anodal stimulation condition with a condition that applied no current at all (no stimulation: 4.27°), the present study compared right anodal stimulation with sham stimulation (sham stimulation: 5.47°). This difference is the reason for the different statistical results with regard to the variable ‘sector width’ in the two studies. In our sham condition, we applied the fade-in and fade-out in such a way that participants felt the same tingling sensation as in the experimental conditions and thus could not distinguish whether it was a real or a control stimulation. In contrast, in the experiment by Bisdorff et al. (1996) it could not be ruled out that the participants knew that it was the control condition if they did not feel the tingling sensation. This might have caused them to react differently and falsify the results than if they had been completely blinded to the experimental conditions. The results obtained in our study also differ from some of the previous studies with regard to the influence of GVS on SPV (see introduction section). One possible reason for the fact that these studies did not observe such an effect could be the small sample sizes used in these earlier studies (Bisdorff et al., 1996; N = 7; Volkening et al., 2014; N = 10). Based on the effect sizes found in the present study, we conducted a power analysis to calculate the minimum sample size needed to find a SPV effect (based on *α* = 0.05, *β* = 0.95 and effect size *f* = 0.34). The necessary sample size turned out to be N = 24 and is therefore considerably larger than the sample sizes actually used in the two aforementioned studies. Further, if not the effect size of the present study but the threshold effect size for a medium effect (*f* = 0.25) is used, the required sample size would be even larger (N = 44).

Studies with non-human primates as well as with humans suggest that the cortical projection of vestibular input comprises several brain structures. These include the intraparietal sulcus, the parietal operculum and the Sylvian fissure with the peri-Sylvian cortex (Frank & Greenlee, 2018; Lopez et al., 2012; zu Eulenburg et al., 2012), including the so-called parieto-insular vestibular cortex (PIVC). The latter was characterized in non-human primates (Grüsser et al., 1990) and probably corresponds to the human posterior insula and retroinsular regions as well as parts of the human inferior parietal lobe (Bense et al., 2001). We know that these structures are not strictly ‘vestibular’ but rather have a multimodal character, representing a significant site for the neural transformation of converging vestibular, auditory, neck proprioceptive and visual input into higher order spatial representations (Karnath & Dieterich, 2006). Neurons of these regions provide us with redundant information about the position and motion of our body in space and thus seem to play an essential role in adjusting body position relative to external space. Interestingly, research has indicated that processing of vestibular input in humans is not symmetrically represented . Specifically, a dominance of the right hemisphere for multisensory (vestibular) cortical areas has been observed in healthy right-handers during vestibular stimulation (Bense et al., 2001; Brandt & Dieterich, 1999; Dieterich et al., 2003; Fasold et al., 2002; Janzen et al., 2008; Schlindwein et al., 2008; Suzuki et al., 2001). Some of these multisensory (vestibular) cortical areas, such as the Sylvian fissure with the associated peri-Sylvian cortex, are part of a brain network responsible for (visuo-)spatial orientation, which is known to have a right-hemispheric dominance as well (Karnath & Dieterich, 2006; Nobre et al., 1997). Likewise, the present study has revealed an asymmetry, in that anodal GVS applied to the right side had a stronger behavioral effect on the SPV compared to GVS applied to the left side. Whether this unexpected asymmetry is related to the known asymmetry of cortical vestibular activation remains an open question for future investigation. In this context, if the brain network in the left hemisphere is indeed less responsive to vestibular signals and spatial orientation, it may require stronger stimulation to evoke a comparable effect at the behavioral level. Therefore, future research could explore whether increasing the stimulation intensity - for example using 2mA - on the left side might elicit a similar effect to that observed on the right side with 1mA in the present study.

Another finding of the present study is the influence of age. In our group of healthy older participants, we found the SPV more biased than in our younger group. This result is in line with literature-based expectations of age-related changes in the vestibular system. In older individuals, the vestibular system shows structural degeneration independent of vestibular diseases (Faraldo-García et al., 2012; Matheson et al., 1999), probably due to a reduced number of otoconia (Walther & Westhofen, 2007) and hair cells (Alberts et al., 2019). Vestibular degeneration can lead to a variety of symptoms, including postural instability (Curthoys, 2000). The decrease in vestibular signals also leads to a dynamic reweighting of different inputs for postural control, such that older individuals tend to rely more heavily on visual and somatosensory input sources (Curthoys, 2000; Peterka & Loughlin, 2004). However, when these other sensory inputs are removed, as was the case in our experiment when participants were blindfolded and well padded in their seats, the brain must rely on the vestibular input to perceive upright body orientation. The present observation that older individuals on average show greater deviations from upright body posture at 0° than younger subjects when measuring the SPV is thus well in line with the expected decrease in sensitivity of the vestibular system with age.

What significance do our results have for possible applications in the therapeutic neurorehabilitative field? Technically, GVS is very easy to apply. Moreover, the stimulation is painless and there are no known undesirable side effects such as vertigo, nausea, etc during or after stimulation as long as the stimulation intensity does not exceed 1-2mA (Balter et al., 2004). Only slight tingling or itching sensations directly under the electrodes have been reported (Balter et al., 2004; Nguyen et al., 2022; Utz et al., 2011). Therefore, there is actually no reason not to use GVS for the treatment of pusher syndrome. However, a limiting factor could be that the numerical effect of GVS on perceived upright body posture seen in the present study was small compared to the ‒ at least in the acute phase of the stroke ‒ large pathologic tilt of ∼18° from which pusher patients suffer (Karnath et al., 2000). In our older sample, the SPV was tilted to the right by 0.9° on average, which is only a small fraction of the pathologic SPV tilt in pusher patients. In fact, the measured values were almost all still within the range considered ‘normal’: a meta-analysis on SPV values in healthy subjects suggested to consider values between -2.87° and 3.11° as non-pathological (Conceição et al., 2018). Only two male participants in our older group had higher values than this, both under right-sided anodal stimulation. However, it remains to be seen whether regular application of GVS over a longer time period may have a lasting therapeutic effect in patients with pusher syndrome. In fact, first applications of GVS in pusher patients have already been carried out, all of which applied anodal GVS to the mastoid located on the side of the lesioned hemisphere. Krewer et al. (2013) found no improvement with GVS in pusher patients. In contrast, Nakamura et al. (2014) observed that GVS could enhance the improvement achieved by physiotherapy. Unfortunately, due to the study design, it was not possible to investigate the effect of GVS alone. Additionally, they applied GVS in a supine and not in a vertical position. In a third study, Babyar et al. (2018) compared the effect of GVS with transcranial direct current stimulation via the PIVC in pusher patients. Unfortunately, the authors did not define a clinically relevant outcome criterion, which makes it difficult to interpret their results in terms of therapeutic use. Further studies are therefore required to clarify the question of a possible effect of anodal GVS on the pathological perception of upright body orientation with respect to gravity in pusher patients.

## Acknowledgements

We would like to thank Kirsti Brandes and Lilli Rötzer for their help with data acquisition.

## Author Contributions

All authors were involved in the conception and design of the study. The recruitment of participants, data collection and analyses were carried out by S.M-W. The first draft of the manuscript was written by S.M-W. and H-O.K. commented on earlier versions of the manuscript. H-O.K. provided supervision and resources and acquired funding. All authors read and approved the final manuscript.

## Data Availability Statement

Upon reasonable request, the corresponding author can provide the data supporting the findings of this study.

## Abbreviations

AL: anode left condition
AR: anode right condition
GVS: galvanic vestibular stimulation
PIVC: parieto-insular vestibular cortex
S: sham condition
SPV: subjective postural vertical
SVV: subjective visual vertical

## References

Alberts, B. B. G. T., Selen, L. P. J., & Medendorp, W. P. (2019). Age-related reweighting of visual and vestibular cues for vertical perception. Journal of Neurophysiology, 121(4), 1279–1288. 10.1152/jn.00481.2018

Babyar, S., Santos, T., Will-Lemos, T., Mazin, S., Edwards, D., & Reding, M. (2018). Sinusoidal transcranial direct current versus galvanic vestibular stimulation for treatment of lateropulsion poststroke. Journal of Stroke and Cerebrovascular Diseases, 27(12), 3621–3625. 10.1016/j.jstrokecerebrovasdis.2018.08.034

Balter, S. G. T., Stokroos, R. J., De Jong, I., Boumans, R., Van De Laar, M., & Kingma, H. (2004). Background on methods of stimulation in galvanic-induced body sway in young healthy adults. Acta Oto-Laryngologica, 124(3), 262–271. 10.1080/00016480310015245

Bense, S., Stephan, T., Yousry, T. A., Brandt, T., & Dieterich, M. (2001). Multisensory Cortical Signal Increases and Decreases During Vestibular Galvanic Stimulation (fMRI). Journal of Neurophysiology, 85(2), 886–899. 10.1152/jn.2001.85.2.886

Bergmann, J., Krewer, C., Selge, C., Müller, F., & Jahn, K. (2016). The subjective postural vertical determined in patients with pusher behavior during standing. Topics in Stroke Rehabilitation, 23(3), 184–190. 10.1080/10749357.2015.1135591

Bisdorff, A., Wolsley, C., Anastasopoulos, D., Bronstein, A., & Gresty, M. (1996). The perception of body verticality (subjective postural vertical) in peripheral and central vestibular disorders. Brain, 119(5), 1523–1534. 10.1093/brain/119.5.1523

Brandt, T., & Dieterich, M. (1999). The Vestibular Cortex: Its Locations, Functions, and Disorders. Annals of the New York Academy of Sciences, 871(1), 293–312. 10.1111/j.1749-6632.1999.tb09193.x

Cohen, B., Yakushin, S., & Holstein, G. (2012a). What Does Galvanic Vestibular Stimulation Actually Activate: Response [Opinion]. Frontiers in Neurology, 3. 10.3389/fneur.2012.00148

Cohen, B., Yakushin, S. B., & Holstein, G. R. (2012b). What Does Galvanic Vestibular Stimulation Actually Activate? [Opinion]. *Frontiers in Neurology*, volume 2 *-* 2011. 10.3389/fneur.2011.00090

Conceição, L. B., Baggio, J. A. O., Mazin, S. C., Edwards, D. J., & Santos, T. E. G. (2018). Normative data for human postural vertical: A systematic review and meta-analysis. PloS One, 13(9), e0204122. 10.1371/journal.pone.0204122

Curthoys, I. S. (2000). Vestibular compensation and substitution. Current Opinion in Neurology, 13(1), 27–30. 10.1097/00019052-200002000-00006

Curthoys, I. S., & MacDougall, H. G. (2012). What Galvanic Vestibular Stimulation Actually Activates [Perspective]. Frontiers in Neurology, Volume 3 - 2012. 10.3389/fneur.2012.00117

Dai, S., Lemaire, C., Piscicelli, C., & Pérennou, D. (2022). Lateropulsion Prevalence After Stroke: A Systematic Review and Meta-analysis. Neurology, 98(15), e1574–e1584. 10.1212/WNL.0000000000200010

Dakin, C. J., & Rosenberg, A. (2018). Chapter 3 - Gravity estimation and verticality perception. In B. L. Day & S. R. Lord (Eds.), Handbook of Clinical Neurology (Vol. 159, pp. 43–59). Elsevier. 10.1016/B978-0-444-63916-5.00003-3

Davies, P. M. (1985). Steps to Follow. A Guide to the Treatment of Adult Hemiplegia (1 ed.). Springer Berlin, Heidelberg.

Day, B. L., & Fitzpatrick, R. C. (2005). The vestibular system. Current Biology, 15(15), R583–R586. 10.1016/j.cub.2005.07.053

Dieterich, M., Bense, S., Lutz, S., Drzezga, A., Stephan, T., Bartenstein, P., & Brandt, T. (2003). Dominance for Vestibular Cortical Function in the Non-dominant Hemisphere. Cerebral Cortex, 13(9), 994–1007. 10.1093/cercor/13.9.994

Faraldo-García, A., Santos-Pérez, S., Crujeiras-Casais, R., Labella-Caballero, T., & Soto-Varela, A. (2012). Influence of age and gender in the sensory analysis of balance control. European Archives of Oto-Rhino-Laryngology, 269(2), 673–677. 10.1007/s00405-011-1707-7

Fasold, O., von Brevern, M., Kuhberg, M., Ploner, C. J., Villringer, A., Lempert, T., & Wenzel, R. (2002). Human Vestibular Cortex as Identified with Caloric Stimulation in Functional Magnetic Resonance Imaging. Neuroimage, 17(3), 1384–1393. 10.1006/nimg.2002.1241

Fitzpatrick, R. C., & Day, B. L. (2004). Probing the human vestibular system with galvanic stimulation. Journal of Applied Physiology, 96(6), 2301–2316. 10.1152/japplphysiol.00008.2004

Frank, S. M., & Greenlee, M. W. (2018). The parieto-insular vestibular cortex in humans: more than a single area? Journal of Neurophysiology, 120(3), 1438–1450. 10.1152/jn.00907.2017

Grüsser, O. J., Pause, M., & Schreiter, U. (1990). Localization and responses of neurones in the parieto-insular vestibular cortex of awake monkeys (Macaca fascicularis). The Journal of Physiology, 430(1), 537–557. 10.1113/jphysiol.1990.sp018306

Janzen, J., Schlindwein, P., Bense, S., Bauermann, T., Vucurevic, G., Stoeter, P., & Dieterich, M. (2008). Neural correlates of hemispheric dominance and ipsilaterality within the vestibularsystem. Neuroimage, 42(4), 1508–1518. 10.1016/j.neuroimage.2008.06.026

Karnath, H.-O. (2007). Pusher syndrome – a frequent but little-known disturbance of body orientation perception. Journal of Neurology, 254(4), 415–424. 10.1007/s00415-006-0341-6

Karnath, H.-O., & Dieterich, M. (2006). Spatial neglect—a vestibular disorder? Brain, 129(2), 293–305. 10.1093/brain/awh698

Karnath, H.-O., Ferber, S., & Dichgans, J. (2000). The origin of contraversive pushing: Evidence for a second graviceptive system in humans. Neurology, 55(9), 1298–1304. 10.1212/WNL.55.9.1298

Kassambara, A. (2023). Pipe-Friendly Framework for Basic Statistical Tests. In (Version 0.7.2) [Package]. https://cran.r-project.org/web/packages/rstatix/rstatix.pdf

Krewer, C., Rieß, K., Bergmann, J., Müller, F., Jahn, K., & Koenig, E. (2013). Immediate effectiveness of single-session therapeutic interventions in pusher behaviour. Gait and Posture, 37(2), 246–250. 10.1016/j.gaitpost.2012.07.014

Latt, L. D., Sparto, P. J., Furman, J. M., & Redfern, M. S. (2003). The steady-state postural response to continuous sinusoidal galvanic vestibular stimulation. Gait and Posture, 18(2), 64–72. 10.1016/S0966-6362(02)00195-9

Lopez, C., Blanke, O., & Mast, F. W. (2012). The human vestibular cortex revealed by coordinate-based activation likelihood estimation meta-analysis. Neuroscience, 212, 159–179. 10.1016/j.neuroscience.2012.03.028

Mars, F., Vercher, J.-L., & Popov, K. (2005). Dissociation between subjective vertical and subjective body orientation elicited by galvanic vestibular stimulation. Brain Research Bulletin, 65(1), 77–86. 10.1016/j.brainresbull.2004.11.012

Matheson, A. J., Darlington, C. L., & Smith, P. F. (1999). Dizziness in the elderly and age-related degeneration of the vestibular system. New Zealand Journal of Psychology, 28(1), 10–16.

Nakamura, J., Kita, Y., Yuda, T., Ikuno, K., Okada, Y., & Shomoto, K. (2014). Effects of galvanic vestibular stimulation combined with physical therapy on pusher behavior in stroke patients: a case series. NeuroRehabilitation, 35(1), 31–37. 10.3233/NRE-141094

Nestmann, S., Karnath, H.-O., Bülthoff, H. H., & de Winkel, K. N. (2020). Changes in the perception of upright body orientation with age. PloS One, 15(5), e0233160. 10.1371/journal.pone.0233160

Nguyen, T. T., Kang, J. J., & Oh, S. Y. (2022). Thresholds for vestibular and cutaneous perception and oculomotor response induced by galvanic vestibular stimulation. Frontiers in Neurology(1664–2295). 10.3389/fneur.2022.955088

Nobre, A. C., Sebestyen, G. N., Gitelman, D. R., Mesulam, M. M., Frackowiak, R. S., & Frith, C. D. (1997). Functional localization of the system for visuospatial attention using positron emission tomography. Brain, 120 ( *Pt 3*), 515–533. 10.1093/brain/120.3.515

Peterka, R. J., & Loughlin, P. J. (2004). Dynamic Regulation of Sensorimotor Integration in Human Postural Control. Journal of Neurophysiology, 91(1), 410–423. 10.1152/jn.00516.2003

Posit Software, P. (2024). RStudio: Integrated Development Environment for R. In (Version 4.4.0) Posit Software, PBC. http://www.posit.co/.

Rouder, J. N., Morey, R. D., Speckman, P. L., & Province, J. M. (2012). Default Bayes factors for ANOVA designs. Journal of Mathematical Psychology, 56(5), 356–374. 10.1016/j.jmp.2012.08.001

Schlindwein, P., Mueller, M., Bauermann, T., Brandt, T., Stoeter, P., & Dieterich, M. (2008). Cortical representation of saccular vestibular stimulation: VEMPs in fMRI. Neuroimage, 39(1), 19–31. 10.1016/j.neuroimage.2007.08.016

Suzuki, M., Kitano, H., Ito, R., Kitanishi, T., Yazawa, Y., Ogawa, T., Shiino, A., & Kitajima, K. (2001). Cortical and subcortical vestibular response to caloric stimulation detected by functional magnetic resonance imaging. Cognitive Brain Research, 12(3), 441–449. 10.1016/S0926-6410(01)00080-5

Tardy-Gervet, M.-F., & Séverac-Cauquil, A. (1998). Effects of galvanic vestibular stimulation on perception of subjective vertical in standing humans. Perceptual and motor skills, 86(3_suppl), 1155-1161. 10.2466/pms.1998.86.3c.1155

Utz, K. S., Korluss, K., Schmidt, L., Rosenthal, A., Oppenländer, K., Keller, I., & Kerkhoff, G. (2011). Minor adverse effects of galvanic vestibular stimulation in persons with stroke and healthy individuals. Brain Injury, 25(11). 10.3109/02699052.2011.607789

Volkening, K., Bergmann, J., Keller, I., Wuehr, M., Müller, F., & Jahn, K. (2014). Verticality perception during and after galvanic vestibular stimulation. Neuroscience Letters, 581, 75–79. 10.1016/j.neulet.2014.08.028

Walther, L. E., & Westhofen, M. (2007). Presbyvertigo-aging of otoconia and vestibular sensory cells. Journal of Vestibular Research, 17(2-3), 89–92.

Zink, R., Bucher, S. F., Weiss, A., Brandt, T., & Dieterich, M. (1998). Effects of galvanic vestibular stimulation on otolithic and semicircular canal eye movements and perceived vertical. Electroencephalography and Clinical Neurophysiology, 107(3), 200–205. 10.1016/S0013-4694(98)00056-X

zu Eulenburg, P., Caspers, S., Roski, C., & Eickhoff, S. B. (2012). Meta-analytical definition and functional connectivity of the human vestibular cortex. Neuroimage, 60(1), 162–169. 10.1016/j.neuroimage.2011.12.032

